## Supplementary Tables Captions for "Transcriptional immune suppression and upregulation of double stranded DNA damage and repair repertoires in ecDNA-containing tumors"

**Table S1.** ecDNA status of 870 TCGA samples across 14 tumor types.

**Table S2.** Highly co-expressed gene clusters identified using multiscale bootstrap resampling.

**Table S3.** Cluster #3 and #74 members.

**Table S4.** Cliff's delta values of 643 CorEx genes in TCGA samples, and in 11 tumor types with at least 10 ecDNA(+) and 10 ecDNA(-) samples each.

**Table S5.** Top-|LFC| genes: 643 most significantly differentially expressed genes based on logarithmic fold changes from a DESeq2 analysis.

**Table S6.** Up-/down-regulated genes in 870 TCGA samples.

**Table S7.** GO biological processes enriched in up-regulated CorEx genes.

**Table S8.** Clustering of GO biological processes enriched in up-regulated CorEx genes into 11 broad categories.

**Table S9.** GO biological processes enriched in Cluster #3 genes.

**Table S10.** Up-regulated CorEx genes unique to biological process categories.

**Table S11.** Hand-curated list of 129 genes involved in DSB repair pathways.

**Table S12.** ecDNA status of 1,440 TCGA samples across 24 tumor types.

**Table S13.** Up-/down-regulated genes in 1,440 TCGA samples.

**Table S14.** GO biological processes enriched in down-regulated CorEx genes.

**Table S15.** Clustering of GO biological processes enriched in down-regulated CorEx genes into seven broad categories.

**Table S16.** Down-regulated CorEx genes in NF-κB signaling (GO:0007249) involved in pro-apoptotic caspase activation.

**Table S17.** Differentially mutated genes in ecDNA(+) and ecDNA(-) samples and their odds ratios.
