## Supplementary Figures for "Transcriptional immune suppression and upregulation of double stranded DNA damage and repair repertoires in ecDNA-containing tumors"

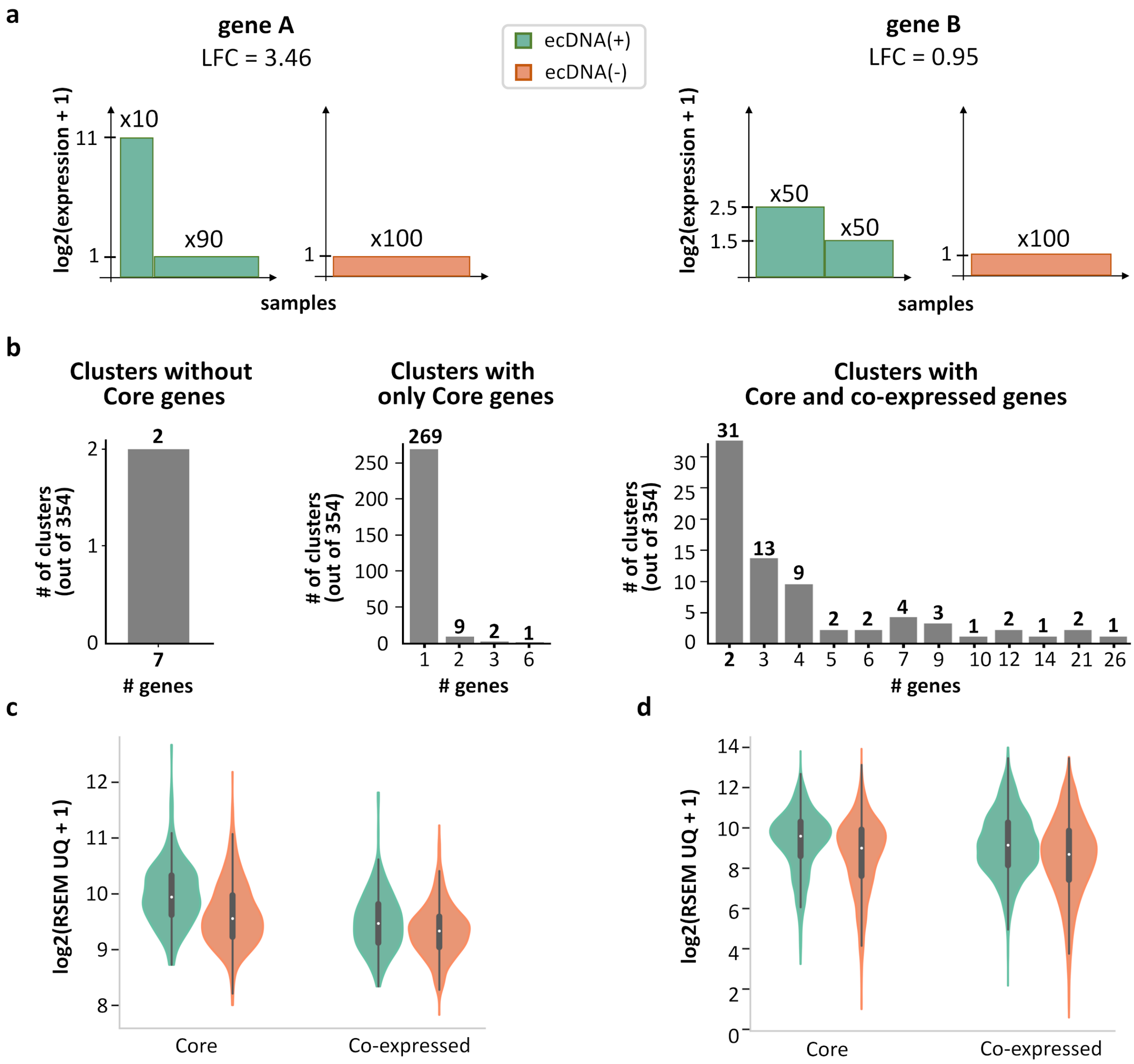


**Figure S1. Cartoon illustration of the rationale for Boruta analysis**. **(a)** A differentially expressed gene (e.g. gene A) may have a higher log-fold change (LFC) value than that of a CorEx gene (e.g. gene B), but may not be persistently over- or under-expressed in ecDNA(+) samples**.** Therefore, no expression value cut-off can be used to separate ecDNA(+) samples from ecDNA(-) samples using gene A expression. In contrast, gene B expression can separate the two classes perfectly. **(b)** Pvclust identified 354 highly co-expressed clusters where members are selected in at least 10 out of 200 Boruta trials as a cluster. Of the 354 clusters, 2 did not contain any Core genes, 281 only contained Core genes, and 71 contained at least 1 Core and 1 co-expressed gene. **(c)** Cluster #74 contains a Core gene, *RAE1*, and a co-expressed gene, *CSTF1*. **(d)** Cluster #3 contains 9 Core genes and 12 co-expressed genes.

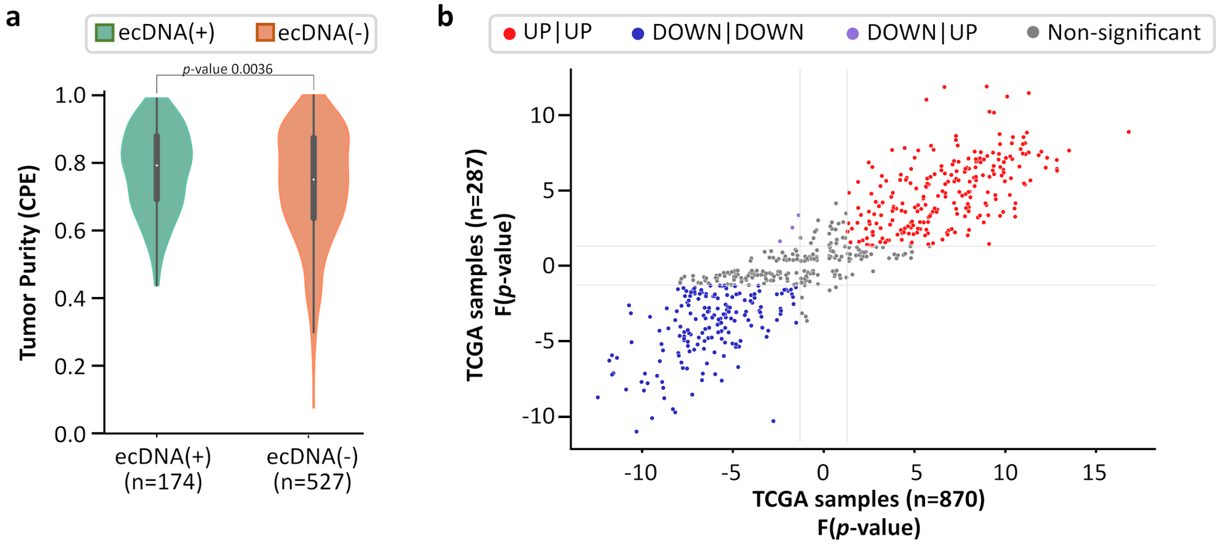


**Figure S2. Impact of tumor purity on CorEx gene expression.** Of the 870 TCGA samples (234 ecDNA(+), 636 ecDNA(-)), 701 samples (174 ecDNA(+), 527 ecDNA(-)) were assigned a consensus measurement of purity estimation (CPE) by Aran et al. **(a)** ecDNA(-) samples have slightly lower purity than ecDNA(+) samples (MWU *p*-value 0.0036). **(b)** For samples with high tumor purity (CPE≥0.8, n=287), the expression directionality of CorEx genes (i.e., based on Mann-Whitney U test) were highly correlated with that of all samples (n=870). Among the 275 genes that had a higher expression in the ecDNA(+) samples of all samples, 243 genes also had a higher expression in the ecDNA(+) samples of high tumor purity samples (red dots). Among the 284 genes that had a lower expression in the ecDNA(+) samples of all samples, 181 genes also had a lower expression in the ecDNA(+) samples of high tumor purity samples (blue dots). Together, 424 of 559 (75.8%) CorEx genes with significant directionality were in agreement between all samples and samples with high tumor purity.

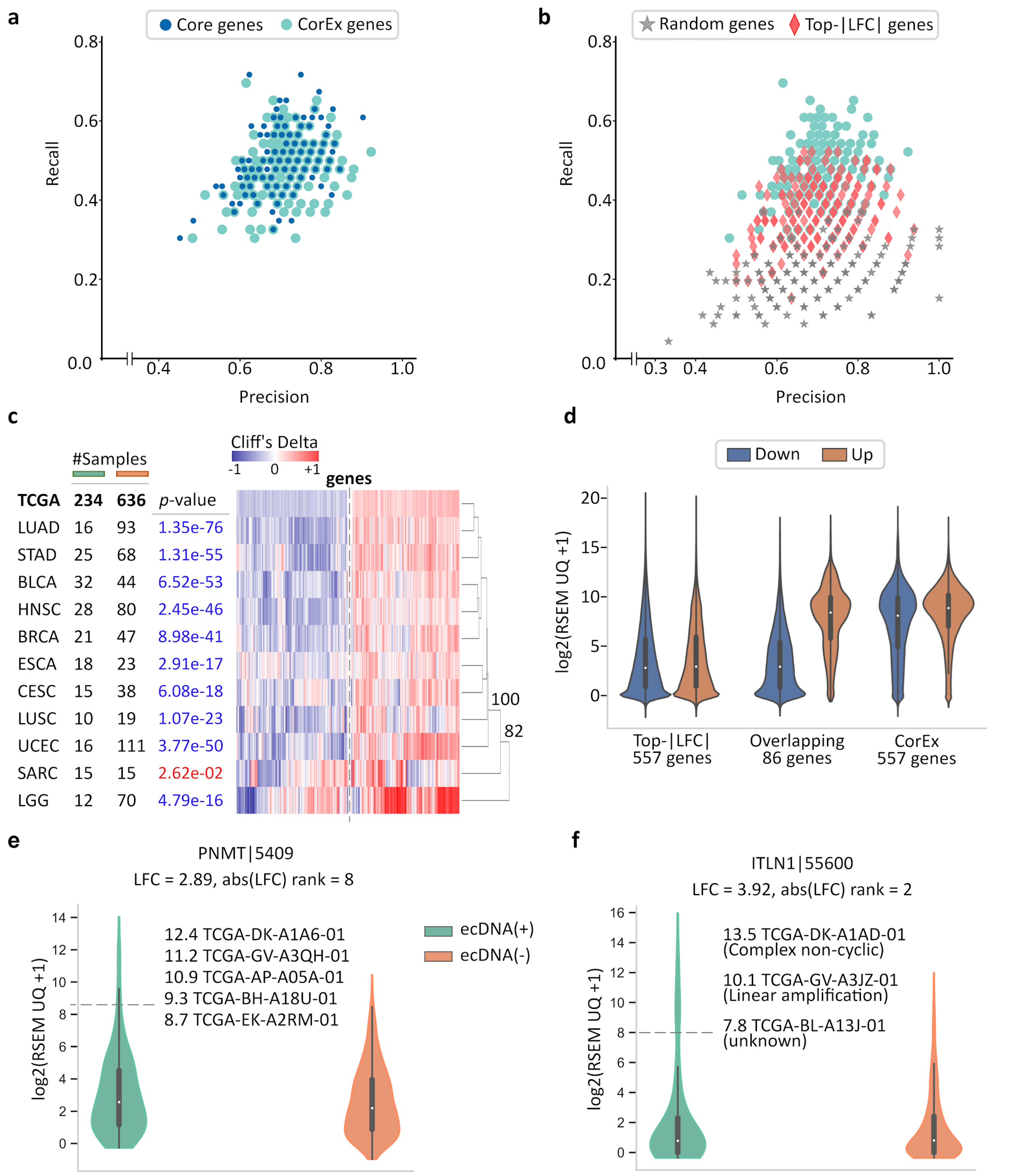


**Figure S3. Extended validation of CorEx genes.** **(a)** Cross-validation experiments validating the predictive value of CorEx genes. Precision denotes the fraction of predicted samples that were truly ecDNA(+). Recall refers to the fraction of ecDNA(+) samples that were predicted correctly. CorEx genes have similar predictive value to Core genes. **(b)** CorEx genes have higher predictive rates compared to 643 randomly selected genes and the top 643 differentially expressed genes based on logarithmic fold changes from a DESeq2 analysis (Top-|LFC| genes). (Continued on the following page.)

**Figure S3 (Continued). (c)** Core genes are also consistently up- or down-regulated in ecDNA(+) samples across tumor types, with the exception of SARC, similar to CorEx genes (Figure 2c). AU *p*-values from multiscale bootstrap resampling are shown at the dendrogram branches. **(d)** 86 CorEx genes are part of the 643 Top-|LFC| gene set. **(e)** *PNMT* normalized gene expression in ecDNA(+) vs. ecDNA(-) samples. **(f)** *ITLN1* normalized gene expression in ecDNA(+) vs. ecDNA(-) samples. In both cases, their up-regulation in ecDNA(+) samples is explained by their presence on a region with somatic, focal copy number amplification.

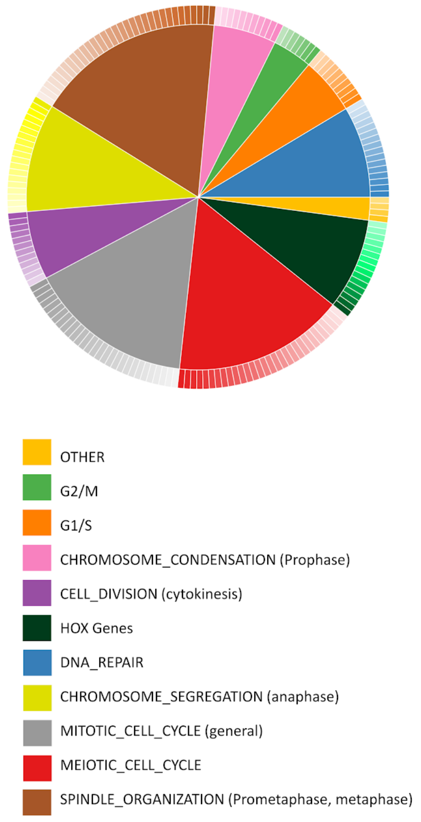


| | padj | GO ID | GO Biological Process Term |
|---|---|---|---|
| 2.1 | 2.28E-02 | GO:1905208 | NEGATIVE_REGULATION_OF_CARDIOCYTE_DIFFERENTIATION |
| | 3.82E-02 | GO:1901563 | RESPONSE_TO_CAMPTOTHECIN |
| | 1.10E-02 | GO:0061351 | NEURAL_PRECURSOR_CELL_PROLIFERATION |
| | 4.42E-02 | GO:0007405 | NEUROBLAST_PROLIFERATION |
| 3.7 | 1.14E-02 | GO:1902750 | NEGATIVE_REGULATION_OF_CELL_CYCLE_G2_M_PHASE_TRANSITION |
| | 2.80E-03 | GO:0044818 | MITOTIC_G2_M_TRANSITION_CHECKPOINT |
| | 3.68E-02 | GO:0007095 | MITOTIC_G2_DNA_DAMAGE_CHECKPOINT_SIGNALING |
| | 1.97E-06 | GO:0044839 | CELL_CYCLE_G2_M_PHASE_TRANSITION |
| | 8.31E-06 | GO:1902749 | REGULATION_OF_CELL_CYCLE_G2_M_PHASE_TRANSITION |
| | 2.47E-02 | GO:1902751 | POSITIVE_REGULATION_OF_CELL_CYCLE_G2_M_PHASE_TRANSITION |
| | 2.47E-02 | GO:0043392 | NEGATIVE_REGULATION_OF_DNA_BINDING |
| 5.4 | 3.11E-04 | GO:0044774 | MITOTIC_DNA_INTEGRITY_CHECKPOINT_SIGNALING |
| | 1.72E-04 | GO:0031570 | DNA_INTEGRITY_CHECKPOINT_SIGNALING |
| | 3.75E-02 | GO:0000076 | DNA_REPLICATION_CHECKPOINT_SIGNALING |
| | 4.65E-02 | GO:0006270 | DNA_REPLICATION_INITIATION |
| | 4.58E-04 | GO:0006260 | DNA_REPLICATION |
| | 1.48E-03 | GO:0006261 | DNA_DEPENDENT_DNA_REPLICATION |
| | 1.00E-02 | GO:0031297 | REPLICATION_FORK_PROCESSING |
| | 1.99E-02 | GO:0045005 | DNA_DEPENDENT_DNA_REPLICATION_MAINTENANCE_OF_FIDELITY |
| | 2.29E-02 | GO:0000723 | TELOMERE_MAINTENANCE |
| | 4.28E-02 | GO:1904358 | POSITIVE_REGULATION_OF_TELOMERE_MAINTENANCE_VIA_TELOMERE_LENGTHENING |
| 5.9 | 4.23E-05 | GO:0030261 | CHROMOSOME_CONDENSATION |
| | 3.42E-04 | GO:0007076 | MITOTIC_CHROMOSOME_CONDENSATION |
| | 3.73E-03 | GO:0032392 | DNA_GEOMETRIC_CHANGE |
| | 4.41E-03 | GO:0071103 | DNA_CONFORMATION_CHANGE |
| | 3.82E-02 | GO:1905821 | POSITIVE_REGULATION_OF_CHROMOSOME_CONDENSATION |
| | 2.57E-04 | GO:0007062 | SISTER_CHROMATID_COHESION |
| | 1.81E-02 | GO:0007064 | MITOTIC_SISTER_CHROMATID_COHESION |
| | 4.06E-13 | GO:0051276 | CHROMOSOME_ORGANIZATION |
| | 8.03E-07 | GO:0033044 | REGULATION_OF_CHROMOSOME_ORGANIZATION |
| | 4.82E-02 | GO:2001252 | POSITIVE_REGULATION_OF_CHROMOSOME_ORGANIZATION |
| | 6.31E-05 | GO:2001251 | NEGATIVE_REGULATION_OF_CHROMOSOME_ORGANIZATION |
| 6.4 | 5.11E-04 | GO:0032465 | REGULATION_OF_CYTOKINESIS |
| | 7.56E-04 | GO:0000910 | CYTOKINESIS |
| | 1.25E-16 | GO:0051301 | CELL_DIVISION |
| | 1.30E-03 | GO:0051302 | REGULATION_OF_CELL_DIVISION |
| | 1.40E-03 | GO:0032467 | POSITIVE_REGULATION_OF_CYTOKINESIS |
| | 3.40E-03 | GO:0000281 | MITOTIC_CYTOKINESIS |
| | 1.16E-02 | GO:0051781 | POSITIVE_REGULATION_OF_CELL_DIVISION |
| | 2.06E-02 | GO:0061640 | CYTOSKELETON_DEPENDENT_CYTOKINESIS |
| | 5.48E-18 | GO:0140014 | MITOTIC_NUCLEAR_DIVISION |
| | 1.09E-05 | GO:0007088 | REGULATION_OF_MITOTIC_NUCLEAR_DIVISION |
| | 3.02E-06 | GO:0051783 | REGULATION_OF_NUCLEAR_DIVISION |
| | 1.11E-06 | GO:0051784 | NEGATIVE_REGULATION_OF_NUCLEAR_DIVISION |
| 8.6 | 4.48E-03 | GO:0009954 | PROXIMAL_DISTAL_PATTERN_FORMATION |
| | 4.63E-02 | GO:0035113 | EMBRYONIC_APPENDAGE_MORPHOGENESIS |
| | 1.21E-02 | GO:0060272 | EMBRYONIC_SKELETAL_JOINT_MORPHOGENESIS |
| | 2.71E-02 | GO:0072498 | EMBRYONIC_SKELETAL_JOINT_DEVELOPMENT |
| | 2.89E-08 | GO:0009952 | ANTERIOR_POSTERIOR_PATTERN_SPECIFICATION |
| | 1.17E-06 | GO:0048706 | EMBRYONIC_SKELETAL_SYSTEM_DEVELOPMENT |
| | 2.08E-06 | GO:0048704 | EMBRYONIC_SKELETAL_SYSTEM_MORPHOGENESIS |
| | 1.22E-05 | GO:0003002 | REGIONALIZATION |
| | 5.68E-04 | GO:0007389 | PATTERN_SPECIFICATION_PROCESS |
| | 1.78E-03 | GO:0009790 | EMBRYO_DEVELOPMENT |
| | 2.03E-03 | GO:0048705 | SKELETAL_SYSTEM_MORPHOGENESIS |
| | 2.14E-03 | GO:0048562 | EMBRYONIC_ORGAN_MORPHOGENESIS |
| | 2.80E-03 | GO:0001501 | SKELETAL_SYSTEM_DEVELOPMENT |
| | 4.14E-03 | GO:0009792 | EMBRYO_DEVELOPMENT_ENDING_IN_BIRTH_OR_EGG_HATCHING |
| | 8.13E-03 | GO:0048568 | EMBRYONIC_ORGAN_DEVELOPMENT |
| | 1.57E-02 | GO:0048598 | EMBRYONIC_MORPHOGENESIS |
| 8.6 | 1.99E-06 | GO:0006281 | DNA_REPAIR |
| | 4.95E-06 | GO:0006974 | CELLULAR_RESPONSE_TO_DNA_DAMAGE_STIMULUS |
| | 4.95E-06 | GO:0006302 | DOUBLE_STRAND_BREAK_REPAIR |
| | 8.48E-03 | GO:2000779 | REGULATION_OF_DOUBLE_STRAND_BREAK_REPAIR |
| | 2.13E-02 | GO:0033554 | CELLULAR_RESPONSE_TO_STRESS |
| | 2.90E-03 | GO:0042770 | SIGNAL_TRANSDUCTION_IN_RESPONSE_TO_DNA_DAMAGE |
| | 9.49E-04 | GO:0051052 | REGULATION_OF_DNA_METABOLIC_PROCESS |
| | 1.00E-03 | GO:0006282 | REGULATION_OF_DNA_REPAIR |
| | 1.25E-03 | GO:2001020 | REGULATION_OF_RESPONSE_TO_DNA_DAMAGE_STIMULUS |
| | 5.32E-03 | GO:2001022 | POSITIVE_REGULATION_OF_RESPONSE_TO_DNA_DAMAGE_STIMULUS |
| | 1.84E-07 | GO:0006259 | DNA_METABOLIC_PROCESS |
| | 8.00E-03 | GO:0051054 | POSITIVE_REGULATION_OF_DNA_METABOLIC_PROCESS |
| | 5.53E-04 | GO:0006310 | DNA_RECOMBINATION |
| | 2.20E-02 | GO:0000018 | REGULATION_OF_DNA_RECOMBINATION |
| | 2.32E-02 | GO:0000725 | RECOMBINATIONAL_REPAIR |
| | 7.82E-03 | GO:0035825 | HOMOLOGOUS_RECOMBINATION |
| 10.2 | 2.68E-05 | GO:0033047 | REGULATION_OF_MITOTIC_SISTER_CHROMATID_SEGREGATION |
| | 1.06E-18 | GO:0000070 | MITOTIC_SISTER_CHROMATID_SEGREGATION |
| | 1.63E-04 | GO:0051256 | MITOTIC_SPINDLE_MIDZONE_ASSEMBLY |
| | 2.61E-18 | GO:0007059 | CHROMOSOME_SEGREGATION |
| | 6.68E-19 | GO:0098813 | NUCLEAR_CHROMOSOME_SEGREGATION |
| | 1.10E-18 | GO:0000819 | SISTER_CHROMATID_SEGREGATION |
| | 1.58E-06 | GO:0051983 | REGULATION_OF_CHROMOSOME_SEGREGATION |
| | 4.05E-07 | GO:0051304 | CHROMOSOME_SEPARATION |
| | 1.08E-05 | GO:1905818 | REGULATION_OF_CHROMOSOME_SEPARATION |
| | 4.30E-04 | GO:0051231 | SPINDLE_ELONGATION |
| | 8.70E-04 | GO:0051255 | SPINDLE_MIDZONE_ASSEMBLY |
| | 1.34E-05 | GO:1902100 | NEGATIVE_REGULATION_OF_METAPHASE_ANAPHASE_TRANSITION_OF_CELL_CYCLE |
| | 4.25E-02 | GO:0031145 | ANAPHASE_PROMOTING_COMPLEX_DEPENDENT_CATABOLIC_PROCESS |
| | 3.82E-02 | GO:1905784 | REGULATION_OF_ANAPHASE_PROMOTING_COMPLEX_DEPENDENT_CATABOLIC_PROCESS |
| | 9.86E-04 | GO:1904666 | REGULATION_OF_UBIQUITIN_PROTEIN_LIGASE_ACTIVITY |
| | 5.96E-03 | GO:0051438 | REGULATION_OF_UBIQUITIN_PROTEIN_TRANSFERASE_ACTIVITY |
| | 1.21E-02 | GO:1904668 | POSITIVE_REGULATION_OF_UBIQUITIN_PROTEIN_LIGASE_ACTIVITY |
| | 2.74E-02 | GO:0051443 | POSITIVE_REGULATION_OF_UBIQUITIN_PROTEIN_TRANSFERASE_ACTIVITY |
| | 3.40E-02 | GO:1903320 | REGULATION_OF_PROTEIN_MODIFICATION_BY_SMALL_PROTEIN_CONJUGATION_OR_REMOVAL |
| 15.5 | 9.23E-21 | GO:0000278 | MITOTIC_CELL_CYCLE |
| | 8.01E-21 | GO:1903047 | MITOTIC_CELL_CYCLE_PROCESS |
| | 3.12E-08 | GO:0007093 | MITOTIC_CELL_CYCLE_CHECKPOINT_SIGNALING |
| | 5.58E-08 | GO:0044772 | MITOTIC_CELL_CYCLE_PHASE_TRANSITION |
| | 8.58E-15 | GO:0007049 | CELL_CYCLE |
| | 1.77E-16 | GO:0022402 | CELL_CYCLE_PROCESS |
| | 1.96E-08 | GO:0044770 | CELL_CYCLE_PHASE_TRANSITION |
| | 4.64E-09 | GO:0000075 | CELL_CYCLE_CHECKPOINT_SIGNALING |
| | 1.19E-11 | GO:0010564 | REGULATION_OF_CELL_CYCLE_PROCESS |
| | 8.93E-10 | GO:0051726 | REGULATION_OF_CELL_CYCLE |
| | 4.27E-09 | GO:1901987 | REGULATION_OF_CELL_CYCLE_PHASE_TRANSITION |
| | 5.70E-09 | GO:0007346 | REGULATION_OF_MITOTIC_CELL_CYCLE |
| | 9.37E-09 | GO:1901990 | REGULATION_OF_MITOTIC_CELL_CYCLE_PHASE_TRANSITION |
| | 1.30E-06 | GO:0090068 | POSITIVE_REGULATION_OF_CELL_CYCLE_PROCESS |
| | 4.69E-06 | GO:0045787 | POSITIVE_REGULATION_OF_CELL_CYCLE |
| | 1.72E-04 | GO:1901992 | POSITIVE_REGULATION_OF_MITOTIC_CELL_CYCLE_PHASE_TRANSITION |
| | 2.50E-04 | GO:0045931 | POSITIVE_REGULATION_OF_MITOTIC_CELL_CYCLE |
| | 8.70E-04 | GO:1901989 | POSITIVE_REGULATION_OF_CELL_CYCLE_PHASE_TRANSITION |
| | 3.07E-07 | GO:0010948 | NEGATIVE_REGULATION_OF_CELL_CYCLE_PROCESS |
| | 4.45E-07 | GO:1901988 | NEGATIVE_REGULATION_OF_CELL_CYCLE_PHASE_TRANSITION |
| | 2.65E-06 | GO:0045930 | NEGATIVE_REGULATION_OF_MITOTIC_CELL_CYCLE |
| | 1.26E-05 | GO:1901991 | NEGATIVE_REGULATION_OF_MITOTIC_CELL_CYCLE_PHASE_TRANSITION |
| | 1.41E-05 | GO:0045786 | NEGATIVE_REGULATION_OF_CELL_CYCLE |
| | 1.01E-03 | GO:1904029 | REGULATION_OF_CYCLIN_DEPENDENT_PROTEIN_KINASE_ACTIVITY |
| | 3.88E-03 | GO:0008284 | POSITIVE_REGULATION_OF_CELL_POPULATION_PROLIFERATION |
| | 1.06E-02 | GO:0051338 | REGULATION_OF_TRANSFERASE_ACTIVITY |
| | 3.18E-02 | GO:0051347 | POSITIVE_REGULATION_OF_TRANSFERASE_ACTIVITY |
| | 3.68E-02 | GO:1904031 | POSITIVE_REGULATION_OF_CYCLIN_DEPENDENT_PROTEIN_KINASE_ACTIVITY |
| | 4.65E-02 | GO:0071900 | REGULATION_OF_PROTEIN_SERINE_THREONINE_KINASE_ACTIVITY |
| 16.0 | 1.23E-02 | GO:0051307 | MEIOTIC_CHROMOSOME_SEPARATION |
| | 1.21E-02 | GO:0045143 | HOMOLOGOUS_CHROMOSOME_SEGREGATION |
| | 2.01E-03 | GO:0016321 | FEMALE_MEIOSIS_CHROMOSOME_SEGREGATION |
| | 5.22E-04 | GO:0051445 | REGULATION_OF_MEIOTIC_CELL_CYCLE |
| | 3.75E-02 | GO:2000241 | REGULATION_OF_REPRODUCTIVE_PROCESS |
| | 1.21E-02 | GO:0061983 | MEIOSIS_II_CELL_CYCLE_PROCESS |
| | 3.82E-02 | GO:0051754 | MEIOTIC_SISTER_CHROMATID_COHESION_CENTROMERIC |
| | 4.91E-03 | GO:0033313 | MEIOTIC_CELL_CYCLE_CHECKPOINT_SIGNALING |
| | 3.22E-02 | GO:0051447 | NEGATIVE_REGULATION_OF_MEIOTIC_CELL_CYCLE |
| | 2.98E-09 | GO:1903046 | MEIOTIC_CELL_CYCLE_PROCESS |
| | 3.54E-08 | GO:0051321 | MEIOTIC_CELL_CYCLE |
| | 4.05E-07 | GO:0061982 | MEIOSIS_I_CELL_CYCLE_PROCESS |
| | 2.59E-06 | GO:0045132 | MEIOTIC_CHROMOSOME_SEGREGATION |
| | 6.06E-04 | GO:0070192 | CHROMOSOME_ORGANIZATION_INVOLVED_IN_MEIOTIC_CELL_CYCLE |
| | 1.62E-07 | GO:0007143 | FEMALE_MEIOTIC_NUCLEAR_DIVISION |
| | 3.19E-05 | GO:0044771 | MEIOTIC_CELL_CYCLE_PHASE_TRANSITION |
| | 4.03E-05 | GO:0008315 | G2_MI_TRANSITION_OF_MEIOTIC_CELL_CYCLE |
| | 4.75E-05 | GO:0007144 | FEMALE_MEIOSIS_I |
| | 1.63E-04 | GO:0090306 | MEIOTIC_SPINDLE_ASSEMBLY |
| | 1.11E-03 | GO:0007056 | SPINDLE_ASSEMBLY_INVOLVED_IN_FEMALE_MEIOSIS |
| | 1.98E-03 | GO:0000212 | MEIOTIC_SPINDLE_ORGANIZATION |
| | 9.50E-03 | GO:1901993 | REGULATION_OF_MEIOTIC_CELL_CYCLE_PHASE_TRANSITION |
| | 3.82E-02 | GO:1901995 | POSITIVE_REGULATION_OF_MEIOTIC_CELL_CYCLE_PHASE_TRANSITION |
| | 1.25E-05 | GO:0007292 | FEMALE_GAMETE_GENERATION |
| | 1.36E-04 | GO:0000003 | REPRODUCTION |
| | 5.00E-03 | GO:0048477 | OOGENESIS |
| | 7.69E-03 | GO:0007276 | GAMETE_GENERATION |
| | 1.57E-02 | GO:0044703 | MULTI_ORGANISM_REPRODUCTIVE_PROCESS |
| | 1.66E-02 | GO:0022412 | CELLULAR_PROCESS_INVOLVED_IN_REPRODUCTION_IN_MULTICELLULAR_ORGANISM |
| | 1.98E-02 | GO:0032504 | MULTICELLULAR_ORGANISM_REPRODUCTION |
| 17.7 | 4.29E-05 | GO:0050000 | CHROMOSOME_LOCALIZATION |
| | 4.75E-05 | GO:0051310 | METAPHASE_PLATE_CONGRESSION |
| | 6.38E-04 | GO:0007080 | MITOTIC_METAPHASE_PLATE_CONGRESSION |
| | 3.91E-02 | GO:0051656 | ESTABLISHMENT_OF_ORGANELLE_LOCALIZATION |
| | 1.91E-02 | GO:0051988 | REGULATION_OF_ATTACHMENT_OF_SPINDLE_MICROTUBULES_TO_KINETOCHORE |
| | 9.50E-03 | GO:0051383 | KINETOCHORE_ORGANIZATION |
| | 2.71E-02 | GO:0051315 | ATTACHMENT_OF_MITOTIC_SPINDLE_MICROTUBULES_TO_KINETOCHORE |
| | 5.33E-06 | GO:0008608 | ATTACHMENT_OF_SPINDLE_MICROTUBULES_TO_KINETOCHORE |
| | 1.78E-03 | GO:0090224 | REGULATION_OF_SPINDLE_ORGANIZATION |
| | 1.40E-02 | GO:0032886 | REGULATION_OF_MICROTUBULE_BASED_PROCESS |
| | 1.57E-02 | GO:0070507 | REGULATION_OF_MICROTUBULE_CYTOSKELETON_ORGANIZATION |
| | 3.97E-03 | GO:0061842 | MICROTUBULE_ORGANIZING_CENTER_LOCALIZATION |
| | 3.08E-02 | GO:0051653 | SPINDLE_LOCALIZATION |
| | 3.68E-02 | GO:0040001 | ESTABLISHMENT_OF_MITOTIC_SPINDLE_LOCALIZATION |
| | 3.42E-04 | GO:1903083 | PROTEIN_LOCALIZATION_TO_CONDENSED_CHROMOSOME |
| | 5.75E-04 | GO:0034502 | PROTEIN_LOCALIZATION_TO_CHROMOSOME |
| | 1.16E-03 | GO:0071459 | PROTEIN_LOCALIZATION_TO_CHROMOSOME_CENTROMERIC_REGION |
| | 1.66E-02 | GO:0031109 | MICROTUBULE_POLYMERIZATION_OR_DEPOLYMERIZATION |
| | 1.69E-02 | GO:0007019 | MICROTUBULE_DEPOLYMERIZATION |
| | 9.57E-13 | GO:1902850 | MICROTUBULE_CYTOSKELETON_ORGANIZATION_INVOLVED_IN_MITOSIS |
| | 2.32E-12 | GO:0007052 | MITOTIC_SPINDLE_ORGANIZATION |
| | 7.05E-12 | GO:0007051 | SPINDLE_ORGANIZATION |
| | 1.35E-08 | GO:0051225 | SPINDLE_ASSEMBLY |
| | 3.07E-07 | GO:0000226 | MICROTUBULE_CYTOSKELETON_ORGANIZATION |
| | 1.17E-06 | GO:0090307 | MITOTIC_SPINDLE_ASSEMBLY |
| | 2.00E-06 | GO:0140694 | NON_MEMBRANE_BOUNDED_ORGANELLE_ASSEMBLY |
| | 2.11E-05 | GO:0007017 | MICROTUBULE_BASED_PROCESS |
| | 2.15E-02 | GO:0007010 | CYTOSKELETON_ORGANIZATION |
| | 2.15E-02 | GO:0070925 | ORGANELLE_ASSEMBLY |
| | 1.08E-03 | GO:0033043 | REGULATION_OF_ORGANELLE_ORGANIZATION |
| | 5.14E-04 | GO:0010639 | NEGATIVE_REGULATION_OF_ORGANELLE_ORGANIZATION |
| | 1.06E-18 | GO:0048285 | ORGANELLE_FISSION |
| | 4.45E-06 | GO:0044784 | METAPHASE_ANAPHASE_TRANSITION_OF_CELL_CYCLE |

**Figure S4. Biological process categories enriched in up-regulated CorEx genes.** (Continued on the following page.)

**Figure S4 (Continued).** 187 enriched GOBP terms represented by 169 up-regulated CorEx genes. The enriched biological processes cluster into 11 categories, which in turn can be grouped into Cell-cycle regulation, Mitotic cell-division, double strand break DNA Damage response, and the *HOX* gene cluster.

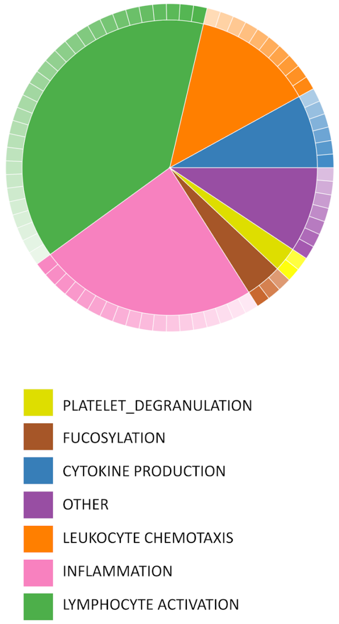


| | padj | GO ID | GO Biological Process Term |
|---|---|---|---|
| 2.67 | 2.28E-02 | GO:0002576 | PLATELET_DEGRANULATION |
| | 4.38E-02 | GO:0090330 | REGULATION_OF_PLATELET_AGGREGATION |
| 4 | 1.19E-02 | GO:0036065 | FUCOSYLATION |
| | 2.28E-02 | GO:0019317 | FUCOSE_CATABOLIC_PROCESS |
| | 4.38E-02 | GO:0009312 | OLIGOSACCHARIDE_BIOSYNTHETIC_PROCESS |
| 8 | 7.17E-03 | GO:0032735 | POSITIVE_REGULATION_OF_INTERLEUKIN_12_PRODUCTION |
| | 7.24E-03 | GO:1903557 | POSITIVE_REGULATION_OF_TUMOR_NECROSIS_FACTOR_SUPERFAMILY_CYTOKINE_PRODUCTION |
| | 7.24E-03 | GO:0071706 | TUMOR_NECROSIS_FACTOR_SUPERFAMILY_CYTOKINE_PRODUCTION |
| | 2.01E-02 | GO:0001816 | CYTOKINE_PRODUCTION |
| | 2.45E-02 | GO:0001819 | POSITIVE_REGULATION_OF_CYTOKINE_PRODUCTION |
| | 3.10E-02 | GO:0032615 | INTERLEUKIN_12_PRODUCTION |
| 9.33 | 1.42E-02 | GO:0007250 | ACTIVATION_OF_NF_KAPPAB_INDUCING_KINASE_ACTIVITY |
| | 3.10E-02 | GO:0035082 | AXONEME_ASSEMBLY |
| | 2.28E-02 | GO:0007249 | I_KAPPAB_KINASE_NF_KAPPAB_SIGNALING |
| | 4.38E-02 | GO:0031294 | LYMPHOCYTE_COSTIMULATION |
| | 4.20E-02 | GO:0030101 | NATURAL_KILLER_CELL_ACTIVATION |
| | 1.19E-02 | GO:0046877 | REGULATION_OF_SALIVA_SECRETION |
| | 3.96E-02 | GO:0046541 | SALIVA_SECRETION |
| 13.33 | 1.76E-03 | GO:0019722 | CALCIUM_MEDIATED_SIGNALING |
| | 3.87E-03 | GO:0060326 | CELL_CHEMOTAXIS |
| | 4.28E-03 | GO:0051480 | REGULATION_OF_CYTOSOLIC_CALCIUM_ION_CONCENTRATION |
| | 4.48E-03 | GO:1990868 | RESPONSE_TO_CHEMOKINE |
| | 7.75E-03 | GO:0050900 | LEUKOCYTE_MIGRATION |
| | 2.78E-02 | GO:0019221 | CYTOKINE_MEDIATED_SIGNALING_PATHWAY |
| | 2.99E-02 | GO:0019932 | SECOND_MESSENGER_MEDIATED_SIGNALING |
| | 3.10E-02 | GO:0030595 | LEUKOCYTE_CHEMOTAXIS |
| | 3.82E-02 | GO:0006873 | CELLULAR_ION_HOMEOSTASIS |
| | 4.38E-02 | GO:0002685 | REGULATION_OF_LEUKOCYTE_MIGRATION |
| 24 | 2.41E-04 | GO:0006955 | IMMUNE_RESPONSE |
| | 7.32E-04 | GO:0006954 | INFLAMMATORY_RESPONSE |
| | 1.38E-03 | GO:0002682 | REGULATION_OF_IMMUNE_SYSTEM_PROCESS |
| | 3.52E-03 | GO:0006952 | DEFENSE_RESPONSE |
| | 4.48E-03 | GO:0034135 | REGULATION_OF_TOLL_LIKE_RECEPTOR_2_SIGNALING_PATHWAY |
| | 7.99E-03 | GO:0034137 | POSITIVE_REGULATION_OF_TOLL_LIKE_RECEPTOR_2_SIGNALING_PATHWAY |
| | 8.35E-03 | GO:0002683 | NEGATIVE_REGULATION_OF_IMMUNE_SYSTEM_PROCESS |
| | 1.01E-02 | GO:0050727 | REGULATION_OF_INFLAMMATORY_RESPONSE |
| | 1.42E-02 | GO:0034134 | TOLL_LIKE_RECEPTOR_2_SIGNALING_PATHWAY |
| | 1.64E-02 | GO:0031347 | REGULATION_OF_DEFENSE_RESPONSE |
| | 1.68E-02 | GO:0050728 | NEGATIVE_REGULATION_OF_INFLAMMATORY_RESPONSE |
| | 2.45E-02 | GO:0050776 | REGULATION_OF_IMMUNE_RESPONSE |
| | 2.49E-02 | GO:0062207 | REGULATION_OF_PATTERN_RECOGNITION_RECEPTOR_SIGNALING_PATHWAY |
| | 3.02E-02 | GO:0031348 | NEGATIVE_REGULATION_OF_DEFENSE_RESPONSE |
| | 3.19E-02 | GO:0045087 | INNATE_IMMUNE_RESPONSE |
| | 3.82E-02 | GO:0050777 | NEGATIVE_REGULATION_OF_IMMUNE_RESPONSE |
| | 4.38E-02 | GO:0002764 | IMMUNE_RESPONSE_REGULATING_SIGNALING_PATHWAY |
| | 4.38E-02 | GO:0034123 | POSITIVE_REGULATION_OF_TOLL_LIKE_RECEPTOR_SIGNALING_PATHWAY |
| 38.67 | 2.00E-05 | GO:0001775 | CELL_ACTIVATION |
| | 2.00E-05 | GO:0046649 | LYMPHOCYTE_ACTIVATION |
| | 2.58E-05 | GO:0042110 | T_CELL_ACTIVATION |
| | 2.41E-04 | GO:0070665 | POSITIVE_REGULATION_OF_LEUKOCYTE_PROLIFERATION |
| | 6.19E-04 | GO:0045058 | T_CELL_SELECTION |
| | 7.24E-04 | GO:0043368 | POSITIVE_T_CELL_SELECTION |
| | 1.38E-03 | GO:0002520 | IMMUNE_SYSTEM_DEVELOPMENT |
| | 1.38E-03 | GO:0002682 | REGULATION_OF_IMMUNE_SYSTEM_PROCESS |
| | 1.45E-03 | GO:0070663 | REGULATION_OF_LEUKOCYTE_PROLIFERATION |
| | 1.76E-03 | GO:0050865 | REGULATION_OF_CELL_ACTIVATION |
| | 2.28E-03 | GO:0070661 | LEUKOCYTE_PROLIFERATION |
| | 4.48E-03 | GO:0046641 | POSITIVE_REGULATION_OF_ALPHA_BETA_T_CELL_PROLIFERATION |
| | 4.48E-03 | GO:0022409 | POSITIVE_REGULATION_OF_CELL_CELL_ADHESION |
| | 7.00E-03 | GO:0042102 | POSITIVE_REGULATION_OF_T_CELL_PROLIFERATION |
| | 7.17E-03 | GO:1903039 | POSITIVE_REGULATION_OF_LEUKOCYTE_CELL_CELL_ADHESION |
| | 7.17E-03 | GO:0050863 | REGULATION_OF_T_CELL_ACTIVATION |
| | 7.24E-03 | GO:0007159 | LEUKOCYTE_CELL_CELL_ADHESION |
| | 7.24E-03 | GO:0051249 | REGULATION_OF_LYMPHOCYTE_ACTIVATION |
| | 9.15E-03 | GO:0050867 | POSITIVE_REGULATION_OF_CELL_ACTIVATION |
| | 2.45E-02 | GO:0050776 | REGULATION_OF_IMMUNE_RESPONSE |
| | 2.78E-02 | GO:0046633 | ALPHA_BETA_T_CELL_PROLIFERATION |
| | 2.78E-02 | GO:0045785 | POSITIVE_REGULATION_OF_CELL_ADHESION |
| | 2.78E-02 | GO:0030217 | T_CELL_DIFFERENTIATION |
| | 3.19E-02 | GO:0022407 | REGULATION_OF_CELL_CELL_ADHESION |
| | 3.19E-02 | GO:0042129 | REGULATION_OF_T_CELL_PROLIFERATION |
| | 3.19E-02 | GO:0042098 | T_CELL_PROLIFERATION |
| | 3.33E-02 | GO:0032753 | POSITIVE_REGULATION_OF_INTERLEUKIN_4_PRODUCTION |
| | 4.20E-02 | GO:1903131 | MONONUCLEAR_CELL_DIFFERENTIATION |
| | 4.20E-02 | GO:0046635 | POSITIVE_REGULATION_OF_ALPHA_BETA_T_CELL_ACTIVATION |

**Figure S5. Biological process categories enriched in down-regulated CorEx genes.** 73 enriched GOBP terms represented by 119 down-regulated CorEx genes. The enriched biological processes can be clustered into 7 categories, 6 of which are related to immune response, suggesting a broad based deregulation.

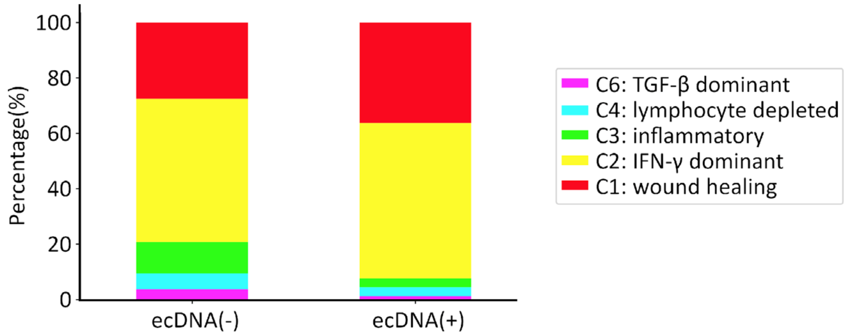


**Figure S6. Tumor microenvironment (TME) subtypes in ecDNA-containing tumors.** TME sub-typing of TCGA samples classified as ecDNA(+) or ecDNA(-) (Amplicon Classifier ver. 0.4.9) based on the immune subtypes provided by Thorsson et al., 2018.

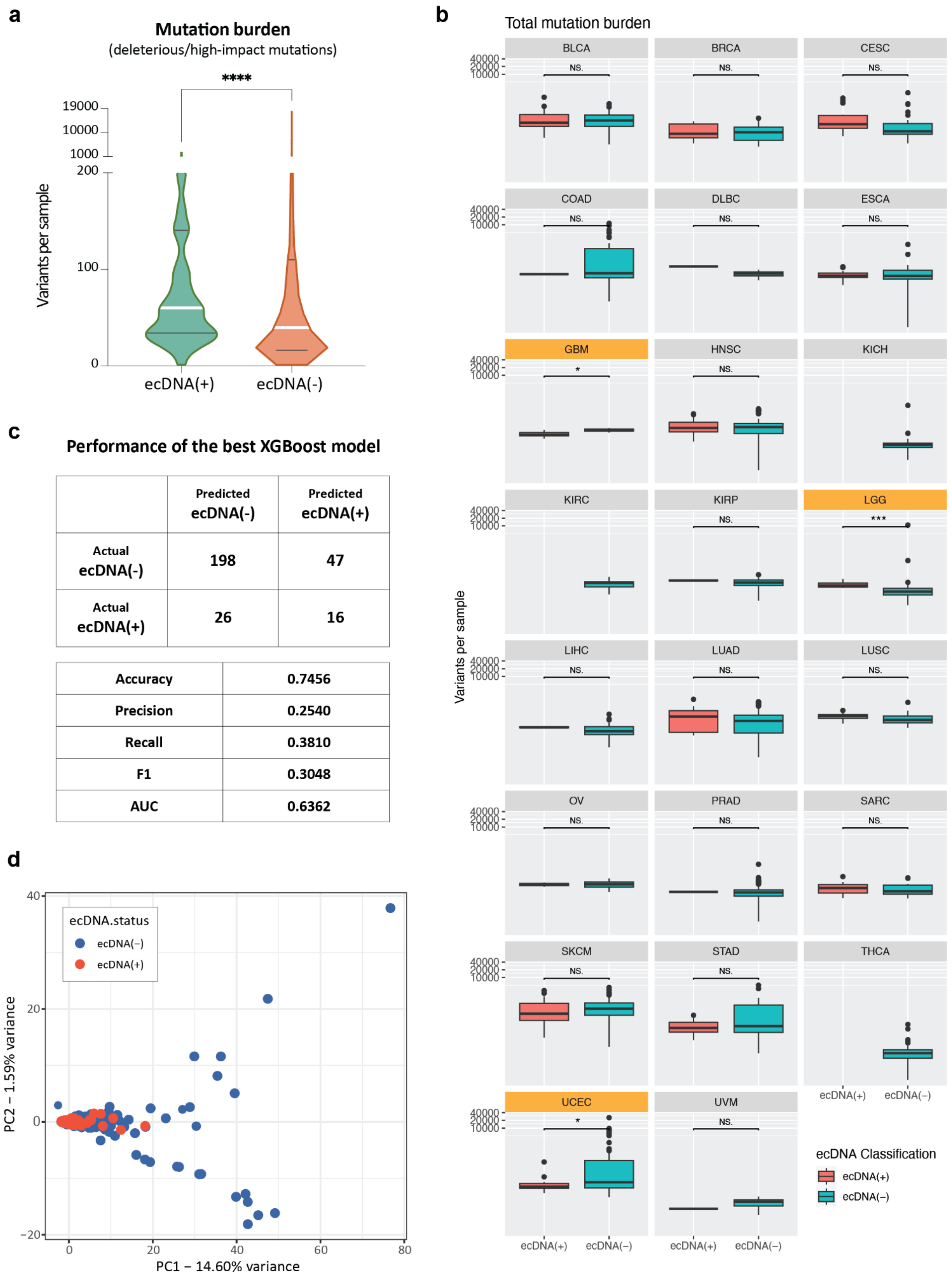


**Figure S7. Extended mutational characteristics of ecDNA-containing tumors. (a)** Mutation burden of ecDNA(+) and ecDNA(-) calculated only with damaging mutations (deleterious SNVs predicted with SIFT or PolyPhen2, frameshift INDELs). (Continued on the following page.)

**Figure S7 (Continued). (b)** Mutation burden by tumor types. Only mutation burdens of GBM, LGG and UCEC were significantly different between ecDNA(+) and ecDNA(-). **(c)** Performance evaluation of the best XGboost model selected by HyperOpt. **(d)** PCA result performed with binary mutational matrix of all mutated genes.

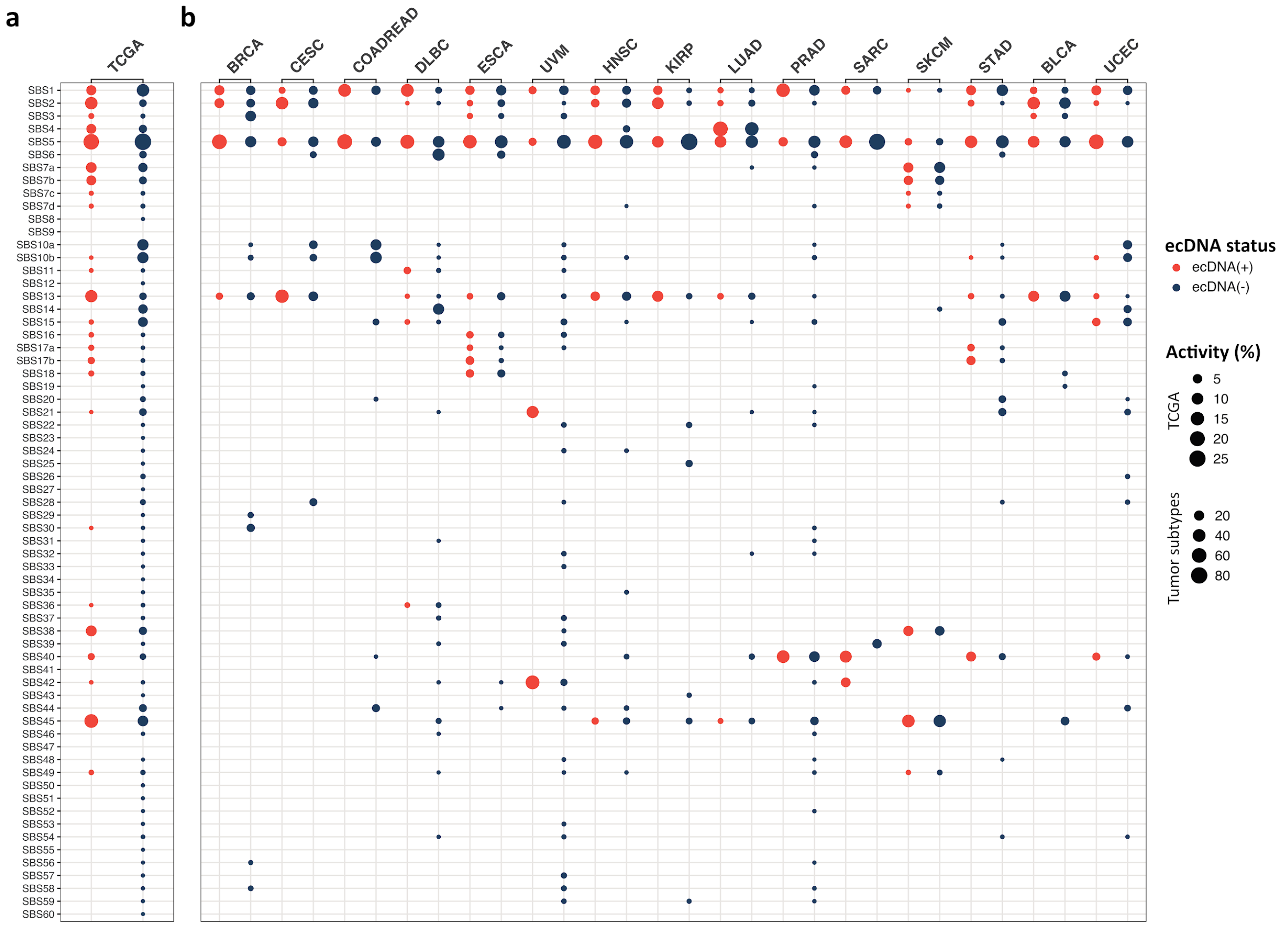


**Figure S8. Single base substitution (SBS) signatures in TCGA samples by ecDNA status.** The activities of SBS signatures 1 to 60 in ecDNA(+) (orange) and ecDNA(-) (blue) samples **(a)** measured across 1,440 TCGA samples and **(b)** separated by tumor types with at least one ecDNA(+) sample each. Activity refers to the estimated relative contribution of a specific mutational signature, and calculated as follows: (number of variants that attributed to a specific signature in a group) / (total number of variants in a group).

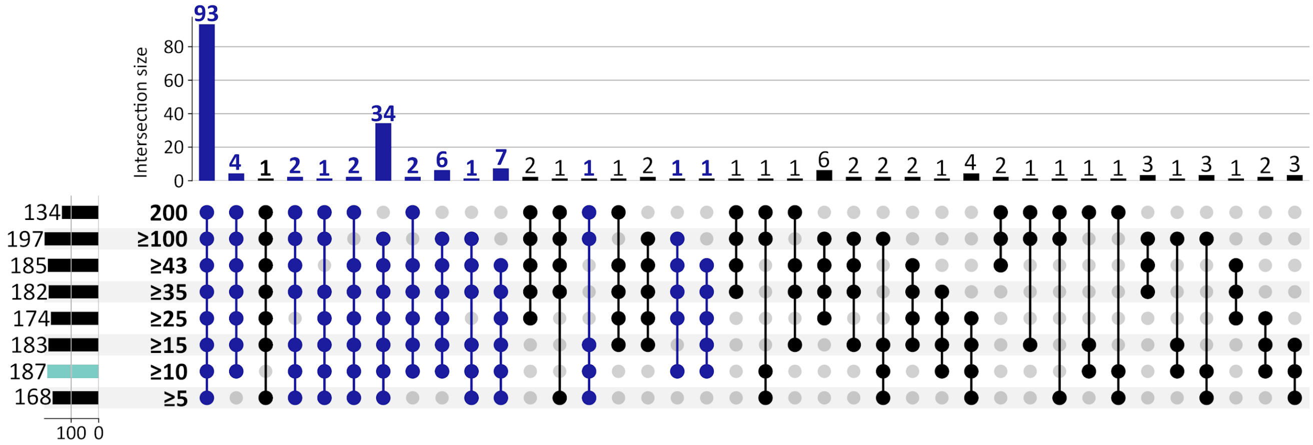


**Figure S9. GO biological processes enriched by up-regulated CorEx genes from 8 selection criteria ranging from 5 to 200 of 200 Boruta trials.** Of the 187 GO terms (turquoise) that were enriched by 262 up-regulated CorEx genes using 10 out of 200 Boruta trials as the selection criteria, 93 terms (49.7%) were enriched for each cut-off criteria, and 155 terms (82.9%) were enriched in at least 5 of the 8 cut-off criteria (dark blue). A minimum degree of 3 was required when using the python UpSet plot function (upsetplot v0.8.0).
